# Back-translating a rodent measure of negative bias into humans: the impact of induced anxiety and unmedicated mood and anxiety disorders

**DOI:** 10.1101/143453

**Authors:** Jessica Aylward, Claire Hales, Emma Robinson, Oliver J Robinson

## Abstract

**Background:** Mood and anxiety disorders are ubiquitous but current treatment options are ineffective for large numbers of sufferers. Moreover, recent years have seen a number of promising pre-clinical interventions fail to translate into clinical efficacy in humans. Improved treatments are unlikely without better animal-human translational pipelines. Here, we directly adapt–i.e. back-translate - a rodent measure of negative affective bias into humans, and explore its relationship with a)pathological mood and anxiety symptoms (study one) and b)transient induced anxiety (study two).

**Method:** Participants who met criteria for mood or anxiety disorder symptomatology according to a face-to-face neuropsychiatric interview were included in the symptomatic group. N = 77(47 asymptomatic; Female = 21; 30 symptomatic; Female = 25) participants completed study one and N = 47 asymptomatic participants (25 female) completed study two. Outcome measures were choice ratios, reaction times and parameters recovered from a computational model of reaction time; the drift diffusion model (DDM).

**Results:** Symptomatic individuals demonstrated increased negative affective bias relative to asymptomatic individuals (proportion high reward = 0.42(SD = 0.14), and 0.53(SD = 0.17), respectively) as well as reduced DDM drift rate (p = 0.004). No significant effects were observed for the within-subjects anxiety-induction in study 2.

**Conclusion:** Humans with pathological anxiety symptoms directly mimic rodents undergoing anxiogenic manipulation. The lack of sensitivity to transient anxiety suggests the paradigm may, moreover, be primarily sensitive to clinically relevant symptoms. Our results establish a direct translational pipeline (and candidate therapeutics screen) from negative affective bias in rodents to pathological mood and anxiety symptoms in humans, and link it to a computational model of reaction time.

## Introduction

Mood and anxiety disorders are extremely prevalent worldwide, with huge psychological, economical and social costs^1^. “Affective biases” which span many domains of cognition, are core features of these disorders^2^. For example, anxious and depressed individuals demonstrate increased sensitivity to aversive stimuli^3^, an attentional bias towards threatening information^2^, and biased interpretation of ambiguous information^4^ (for a review see^5^). These biases both precipitate the onset of disorders and contribute to their maintenance(^5 – 7^). Targeting these biases is therefore a key goal of treatment development.

Unfortunately, for a sizeable number of individuals, current treatments do not lead to clinical improvement^8,9^. Recent years have moreover seen a number of high-profile failures in drug development^10,11^. Among the reasons for this^8,9^, is that some pre-clinical animal tests do not adequately translate the human behaviour they are designed to model^10 – 12^. Indeed there are no tasks that are identical across species; some prominent examples - the forced swim test^13^, or tail suspension test^14^ – do not have clear human analogues. We argue, therefore, that developing *identical* paradigms across humans and animal models will help reduce pre-clinical to clinical translation failure. Instead of *scaling-back* paradigms developed in humans into animals, the present paper takes a paradigm developed within the constraints of an animal model, and directly *‘back-translates’* it for human use.

Specifically, we back-translate a rodent model of affective bias into humans. In the animal task^15^, rats learn to correctly respond to high or low frequency tones, which are associated (100%) with high or low rewards (food pellets). In the test-phase they also respond to an ambiguous mid-tone randomly reinforced with both outcomes. The proportion of low reward responses made to the ambiguous tone represents the degree of negative affective bias. Rats administered an anxiogenic drug or subjected to chronic stress (repeated restraint stress and social isolation)^15^ display increased negative affective bias in choice behaviour. No significant behavioural effect is observed for rats undergoing acute stress (restraint) manipulation.

Here, we explored the impact of two types of anxiety on a human version of this task: a) pathological anxiety in mood and anxiety disorders, and b) acute stress induced using threat of unpredictable shock. The latter stress induction is a well-validated and reliable technique, also back-translated from animal models^16,17^. Critically, it allows the interaction between cognition and anxiety to be explored within-subjects. It elicits ‘adaptive anxiety’ responses such as response inhibition and harm avoidance^17–19^ as well as ‘negative bias’^16,20,21^ in healthy individuals. A related, albeit more complex, version of the present task has previously been tested in healthy participants. Participants responded to a tone paired with reward (to obtain money) and a tone paired with punishment (to avoid punishment). In a test phase participants made more avoidance responses to an ambiguous tone, demonstrating a bias towards avoiding punishment – i.e. a positive bias ^22^. Notably, positive bias responding was correlated negatively with self-reported *state* anxiety level. As such, we predicted that on our novel, directly back-translated rodent task, induced and pathological anxiety would be associated with reduced positive bias – i.e. a negative affective bias.

Computational models can make specific predictions about the underlying mechanisms that drive behaviour and enable a more fine-grained view of decision-making and how it changes in pathological states^23^. One such model – the drift diffusion model – has been applied to rodent data on this task^15^. This model parameterises decision-making as a process of noisy accumulation of evidence^24^. Negative bias following acute pharmacological manipulation and chronic stress in rats was accompanied by increased ‘boundary separation’ parameters (more information required in order to reach a decision), whereas reduced ‘drift rate’ (rate of information accumulation) parameters were seen following the pharmacological manipulation. In this paper we applied both the EZ drift model^25^ - a pared down version of the drift diffusion model^26^ as well as a Bayesian hierarchical drift diffusion model^27^ to our human data.

We therefore tested two predictions. Firstly, considering the well-documented biases in pathological anxiety^28^ and prior work with related tasks^22^, we predicted that individuals with mood and anxiety disorders, relative to the asymptomatic group, would demonstrate increased negative affective bias in this task. Secondly, as induced anxiety instantiates biases across cognition^20^, we predicted that in asymptomatic individuals, threat of shock would also instantiate a negative affective bias. In both cases, we predicted that negative bias in choice behaviour would be associated with alterations to drift diffusion parameters.

## Method

### Participants

Participants were recruited using internet advertisements and via subject databases held at University College London. The only group difference in recruitment was the wording of the advertisements; asymptomatic healthy participants replied to advertisements asking for participants with no psychiatric symptoms; whilst participants with low mood and/or anxiety symptoms replied to advertisements asking for participants who self-defined as experiencing persistent low mood/anxiety symptoms.

A total of 77 participants were included in study 1: 47 asymptomatic participants (Mean age = 28.83, SD = 10.52; 25 female) and 30 (N = 31 originally, but one excluded as they failed to follow task instructions), unmedicated participants with low mood and/or anxiety symptoms (mean age = 28.93, SD = 10.92; 21 Female). A total of 47 asymptomatic participants were included in study 2 (Mean age = 28.96, SD = 10.45; 25 female; 46 overlap with study 1). The neutral version of the task (study 1) was always completed first to ensure consistency with the symptomatic group (who did not complete the stress version). Participants could be aged between 18 – 65 years.

### Symptomatic group details

As depressive and anxiety symptoms are highly comorbid and may not have distinct underlying causes, we include a mixed sample in our symptomatic group (see Table 1 and Supplement). Following an initial screening process, participants who met criteria for mood or anxiety disorder symptomatology according to a face-to-face Mini International Neuropsychiatric Interview (M.I.N.I.^29^) were included in the symptomatic group, those who did not meet any (past/present) criteria according to the M.I.N.I. were included in the asymptomatic group. The State-Trait Anxiety Inventory (STAI^30^) was also collected, as well as additional measures (see Table 1 for full details). Exclusion criteria are listed in the supplement.

**Table 1:** Demographic Information. HC = asymptomatic Healthy Control; ANX = symptomatic individual; STAI = State-Trait Anxiety Inventory; BDI = Beck depression Inventory; Ham-D = Hamilton depression inventory; Ravens’s = Raven’s Progressive Matrices

| **Demographic** | **HC** | **ANX** |
| --- | --- | --- |
| Age | 28.83 (10.52) | 28.93 (10.92) |
| Female | 55.32% | 70% |
| Caucasian | 48% | 56.67% |
| Education | 16.68 (2.23) | 14.37 (5.42) |
| **Clinical** |  |  |
| STAI State | 29.94 (8.06) | 51.83 (9.82) |
| STAI Trait | 33.72 (10.12) | 59.43 (10.05) |
| BDI | 3.30 (3.78) | 25.10 (10.36) |
| Ham-D | - | 16.14 (5.52) |
| Raven's | 8.96 (2.13) | 8.21 (2.74) |

## Procedure

Participants provided written informed consent to take part (UCL ethics reference: 6198/001 or 1764/001). They completed a task coded using the Cogent (Wellcome Trust Centre for Neuroimaging and Institute of Cognitive Neuroscience, UCL, London, UK) toolbox for Matlab (2014b, The MathWorks, Inc., Natick, MA, United States). Scripts available here: 10.6084/m9.figshare.4868303.

### Acquisition task

A task schematic is presented in figure 1. During the acquisition block, participants heard high (1000Hz) and low tones (500Hz), these frequencies were lower than the rat task to account for cross-species differences in hearing. The two tones were associated with different reward values (tone/reward pairings were counterbalanced across participants). They were instructed to learn to make correct key presses following each tone (“z” or “m” key on a laptop keyboard) and informed that correct responses would be rewarded. They were told that they should try and maximise earnings. 10 low and 10 high tones, randomly presented, were played during the practice block. A tone was played for 1000ms followed by an inter stimulus interval of 750ms. A white fixation cross appeared in the middle of the screen during this time. Participants could make their response from the onset of the tone presentation. Following the key press feedback was provided. “*Correct, Win £1*” appeared for 750ms following a correct response to the low reward tone (low/high frequency counterbalanced). “*Correct, Win £4*” appeared for 750ms following a correct response to the high reward tone (low/high frequency, counterbalanced). “*Timeout for incorrect response*” appeared for 3250ms following an incorrect or slow response. The acquisition block enabled participants to understand the key/tone pairings (counterbalanced across participants). The practice block could last between 50-100 seconds.

**Figure 1.**
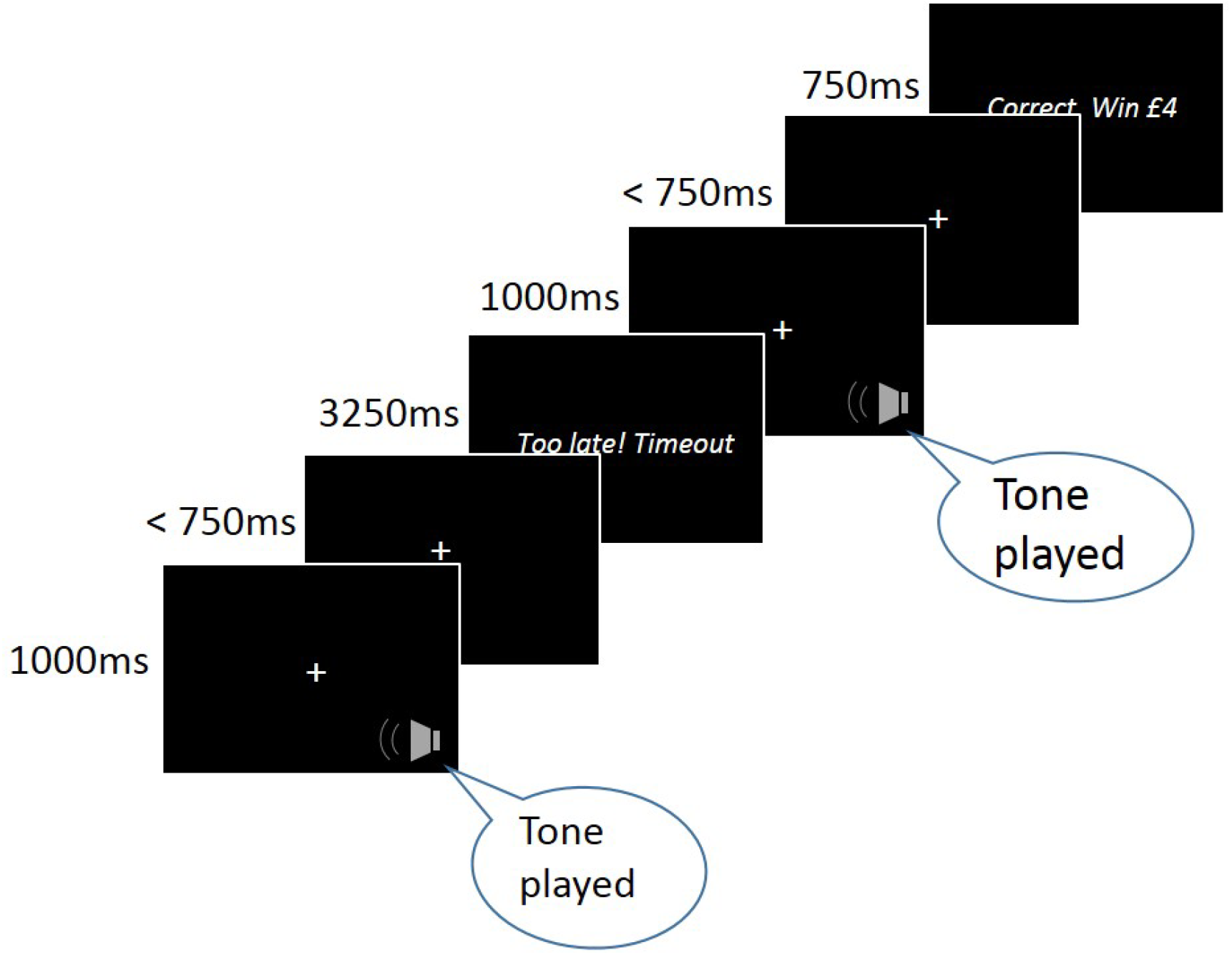
Participants were required to make a key press (“z” or “m” key) following a tone played for 1000ms. After making their response, participants received feedback on their performance. Correct responses saw feedback appear on the screen for 750ms, whilst incorrect responses, or responses made outside the 750ms window, saw feedback appear on the screen for 3250ms. The task consisted of 120 trials, during which 40 low, mid-tone and high tones were presented.

### Testing phase

The tone/reward pairings remained the same as in acquisition but the participants were also presented with a mid-point, ambiguous tone (750Hz) which fell directly in between the low and high tones. Participants were informed that they might hear other tones and that if the tone was unclear, that they should make a key press that corresponded to the closest tone. For half of the trials this mid-tone was associated with a high reward outcome, and for the other half of the trials it was associated with a low reward outcome. As in the practice block, a tone was played for 1000ms, followed by an interstimulus interval of 750ms. Participants made their response as quickly as possible following the tone presentation. Following correct responses the feedback was presented on the screen for 750ms, whilst following incorrect or slow responses *“Timeout for incorrect response”* was presented on the screen for 3250ms.

### Study 1 Symptomatic group vs asymptomatic controls

#### Details

The main task consisted of 120 trials (40 low/mid/high tones, randomly presented). The main task could therefore last between 300–600 seconds.

### Study 2: Induced anxiety version

#### Shock work-up

A Digitimer DS5 Constant Current Stimulator (Digitimer Ltd., Welwyn Garden City, UK) delivered the shocks, via two electrodes attached to the participant’s non-dominant wrist. The shock intensity was increased until the subjective rating was “unpleasant, but not painful”^31^.

#### Stimuli details

A task schematic is presented in figure 2. The task was performed under instructed threat and safe conditions in the same manner as^17^. Participants were told that they would be at risk of an unpredictable shock (independent of their behavioural response), during a threat block (red background). Participants were told that they would be free from shock during a safe block (blue background). Colours were not counterbalanced as prior work has shown this effect to be independent of background colour^32,33^. Each block (total = 4) consisted of 60 randomly presented trials (20 low/mid/high tones; total = 240). The maintenance task could therefore last between 600 – 1200 seconds. Participants either received a shock in the first threat block (post-threat-trial = 45), in the second threat block (post-threat-trial = 96) or at both of these times (randomised across participants). As a manipulation check, participants retrospectively rated their anxiety (out of 10) under threat and safe.

**Figure 2.**
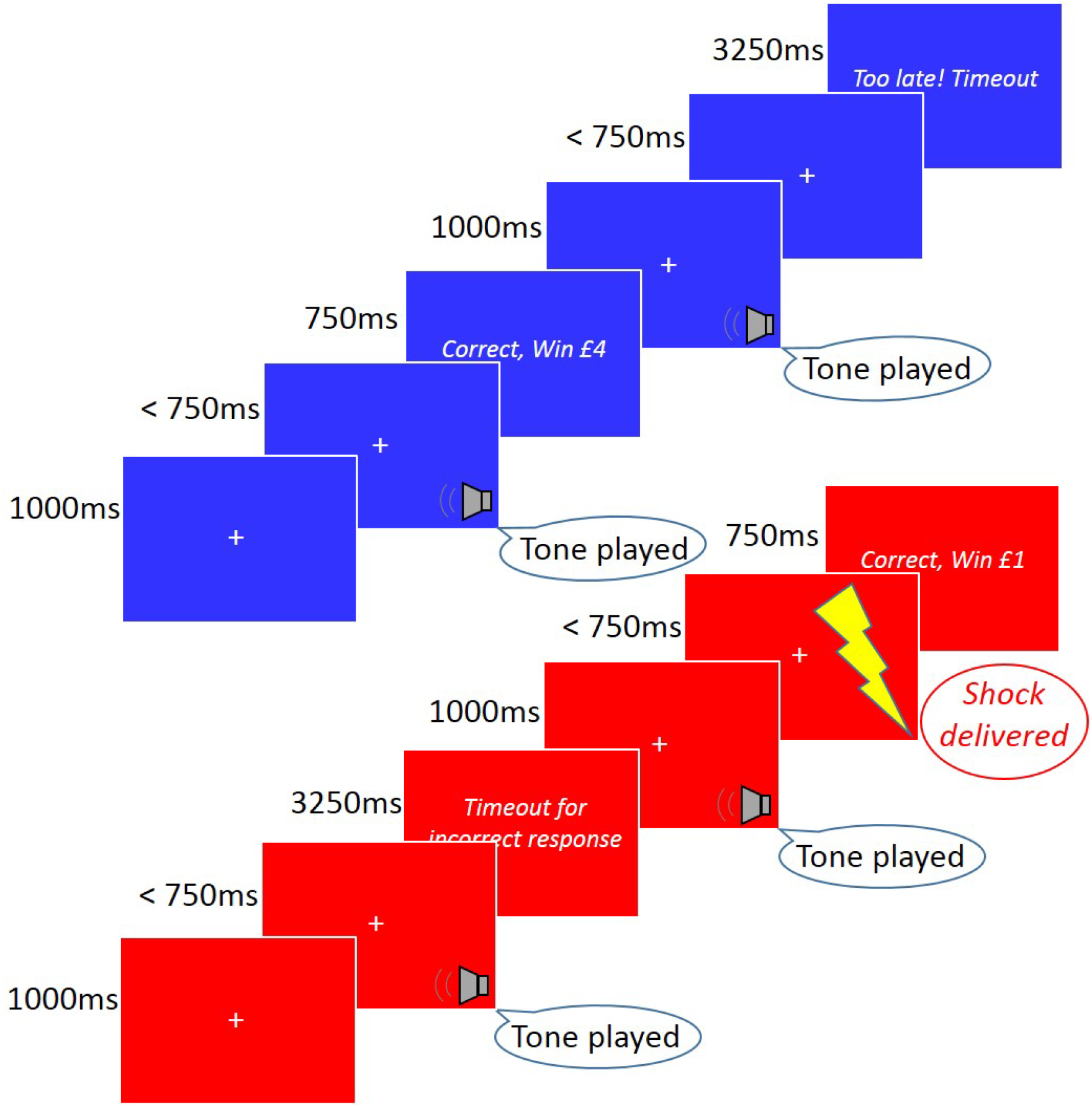
Participants were required to make a key press (“z”/”m”) following a tone played for 1000ms. After making their response, participants received feedback on their performance. Feedback for correct responses lasted 750ms, whilst feedback for incorrect (or slower than 750ms) responses lasted 3250ms. During the safe condition, in which the background was blue, participants were not at risk of shock. During the threat condition, in which the background was red, participants were at risk of unpredictable electric shock.

**Figure 3.**
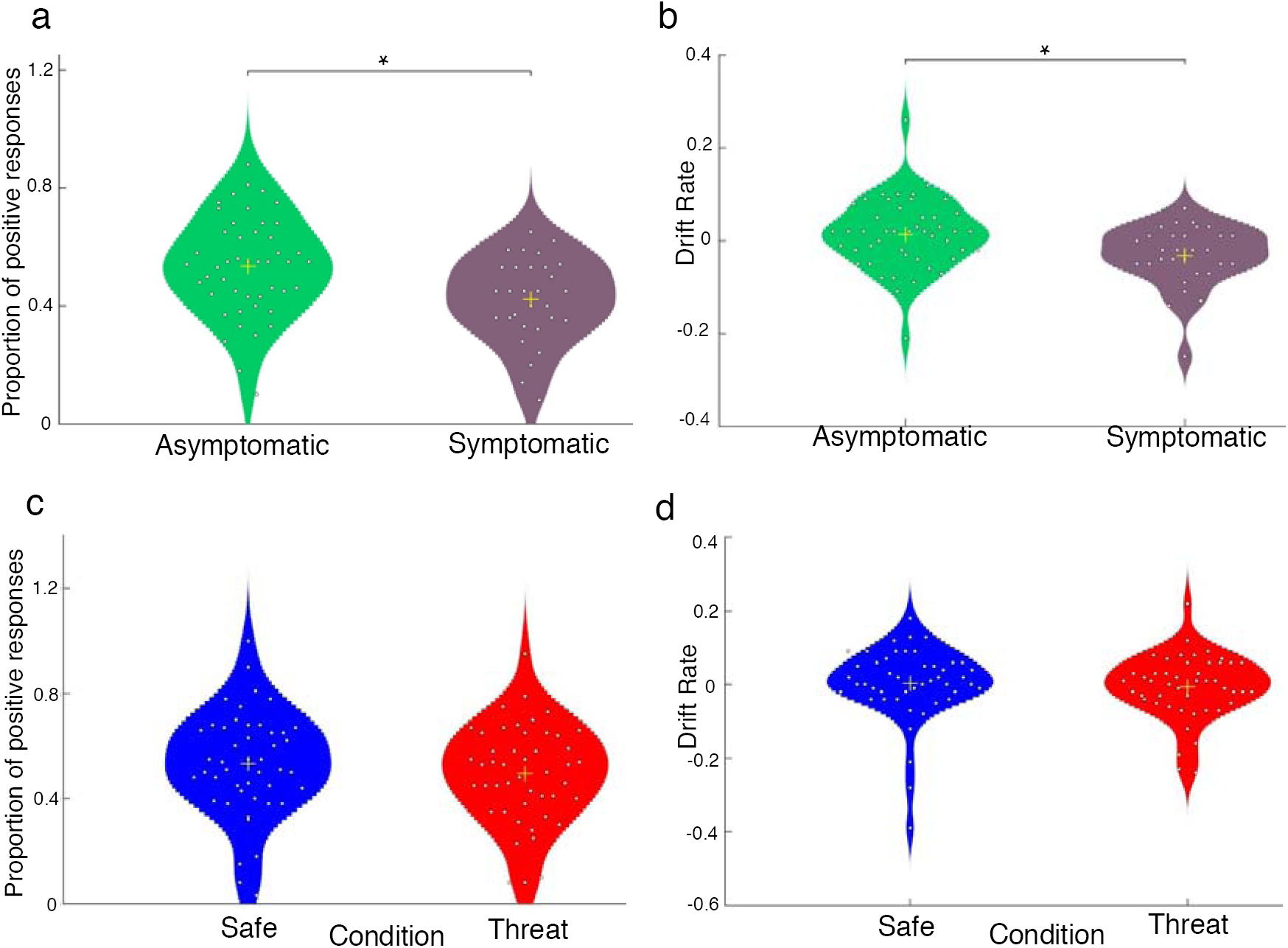
The impact of pathological and induced anxiety on task performance. Violin plots of the proportion of positive responses made to ambiguous tone and EZDM ‘drift rate’-the rate of accumulation of evidence to classify a tone as high reward (shaded area represents a smoothed histogram; yellow cross represents the mean; each circle represents an individual). **a**) Symptomatic individuals had more negative bias (*p* = 0.003, BF_10_ = 12.51) and **b**) a more negative drift rate towards classifying the mid-tone as high reward (*p* = 0.008, BF_10 =_ 5.22). However, there was **c)** no significant difference in affective bias following induced anxiety (*p* = 0.06, BF_10_ = 0.863) **d**) and no significant difference in drift rate across conditions (*p*>0.125, BF_10_<1).

## Statistical analyses

Reaction time (RT) and bias measures (data available here: 10.6084/m9.figshare.4868303) were analysed using SPSS Version 22 (IBM Crop, Armonk, NY). For all analyses, p = 0.05, was considered significant. Affective bias (percentage of ambiguous tones classified as high reward) was calculated by dividing the number of ‘high reward’ responses made to the mid-tone by the total number of key presses made to the mid-tone. RT to respond to the mid-tone was normally distributed and was analysed using independent and paired sample t-tests for study 1 and 2 respectively.

Bayesian statistics were run (JASP, version 0.7^34^), employing the default prior. The Bayesian approach considers the likelihood of the data if the alternative hypothesis is true versus if the null hypothesis is true, allowing for inferences to be made about which model best explains the data. Bayesian ANOVAs and t-tests were used to generate BF_10_ factors which provided evidence for a model of interest relative to a null model. A model with a BF_10_ >1 signifies that model is better at explaining the data relative to the null model, and vice versa for BF_10_ <1. To interpret the magnitude differences between models the following labels were assigned to BF_10_: anecdotal (1-3), substantial (3-10), strong (10-30) decisive (>100)^35^.

Mean RT, variance and proportion of positive responses to the mid-tone were also fed into the EZ drift diffusion model (script available here: 10.6084/m9.figshare.4868303). The parameters of interest were: boundary separation (a), drift rate (v) and non-decision time (t). These refer to the amount of information required before a response can be made (a), the rate at which this information is accumulated (v) and the proportion of the RT that is not accounted for by evidence accumulation (t).

Finally, EZ-DM analyses were supplemented by full hierarchical Bayesian model comparison using the Hierarchical Bayesian estimation of the Drift-Diffusion Model in Python (HDDM) toolbox^27^. The modelled parameters were identical to the above, but this approach also enabled the inclusion of a bias parameter (z), which denotes the starting point between the boundaries. The data for all trials was included in this analysis (stratified into ambiguous mid tone and unambiguous high/low reward trial types) and parameters fit using an Markov Chain Monte Carlo (MCMC) sampling approach implemented using PyMC^36^ (2000 MCMC samples with a burn-in of 20 samples; all winning models obtained Gelman-Rubin statistics ∽1). The influence of adding and subtracting parameters was examined by comparing deviance information criterion (DIC) scores across models. The most extreme 5% of RTs was excluded from all model fitting to account for lapses and facilitate model fitting. Follow-up analysis on recovered parameters was run in a comparable manner to the EZ diffusion analysis and supplemented with a full Bayesian model comparison approach in which the impact of including group or condition in the hierarchical model was tested, and the posteriors of parameters that depended on additional hierarchy plotted for models achieving or exceeding parity of model fits with the basic model. The winning models showed good parameter recovery on posterior predictive checks

Correlation analyses were also run to investigate correlations between STAI trait anxiety scores, affective bias and drift rate.

## Results

### Study 1

#### Choice behaviour

High and low tone accuracy was high (Table 2) and comparable across groups (*t*_*(75)*_ = 0.96, p = 0.338, *d* = 0.22, and *t*_*(75)*_ = 0.28, p = 0.78, *d =*−0.06, respectively). However, there was a significant effect of group on mid-tone choice (*t*_(75)_ = 3.08, p = 0.003, *d =* 0.732, See Fig.3). The symptomatic group were less likely to associate the mid-tone with high reward compared to the asymptomatic group. Bayesian analysis provided strong evidence for a significant difference in affective bias between groups (BF_10 =_ 12.51).

**Table 2:** Average choice, accuracy and reaction time (ms) to all tones in study 1

| **Asymptomatic** | **Accuracy (Standard Deviation)** |
| --- | --- |
| Low Tone | 0.98 (0.05) |
| High Tone | 0.93 (0.083) |
| **Symptomatic** | **Accuracy** |
| Low Tone | 0.97 (0.039) |
| High Tone | 0.95 (0.069) |
| **Group** | **Proportion high reward responses to mid-tone** |
| Asymptomatic | 0.53 (0.17) |
| Symptomatic | 0.42 (0.14) |
| **Asymptomatic** | **Reaction Time** |
| Low Tone | 819.51 (212.00) |
| Mid-tone | 942.41 (181.78) |
| High Tone | 757.33 (228.47) |
| **Symptomatic** | **Reaction Time** |
| Low Tone | 763.17 (197.78) |
| Mid-tone | 894.54 (203.57) |
| High Tone | 694.56 (194.32) |

#### Reaction time

See Table 2 for average RT to all tone types across group. Time to respond to the mid-tone did not differ across groups (*t*_(75)_ = 1.08, p = 0.29, *d =* 0.248). Bayesian analysis favoured the null model (BF_10_ = 0.40).

### DDM

#### EZ-DM

Despite comparable overall RTs there was a significant difference in drift rate between groups (*t*_(75) =_ 2.70, p = 0.008); but not boundary separation (*t*_(75) =_−0.79, p = 0.43) or non-decision time (*t*_(75) =_ 1.3, p = 0.96). The symptomatic group had a slower drift rate towards making a positive choice to the mid-tone (asymptomatic mean = 0.013, SD = 0.075, symptomatic mean =−0.032, SD = 0.066; see Fig.3). Bayesian analysis provided substantial evidence for a significant difference between groups in drift rate (BF_10 =_ 5.22; all other BF_10_<0.31).

#### HDDM

A wide model search was completed (see supplement) across a range of parameters and within-subject factors. The 3 best models are presented in figure 4a. The winning model comprised a model with drift rate, boundary separation, bias and non-decision time parameters (fitted separately across ambiguous mid tone and unambiguous trial types). As with the EZ-DM model, parameters extracted from this winning model demonstrated significant difference in ambiguous mid-tone drift rate between groups (t_(75)_ = 3.0, p = 0.004); but not boundary separation (t_(75)_ =−1.2, p = 0.22), non-decision time (t_(75)_ = 1.4, p = 0.15) or bias (t_(75)_ =−1.4, p = 0.89). The winning model parameters showed a tight correspondence (all r>0.8, p<0.001) with the EZ-DM parameters (see drift rate; figure 4b). However, one advantage of the full hierarchical approach is that we can include group in the model fitting procedure. This approach revealed a winning model (of equivalent fit to the model fit across groups) where the drift rate parameter alone is separated by group. Posterior distributions demonstrate that this is because v on mid tones is lower in patients relative to controls (figure 4d). In short, the full hierarchical model is consistent with the basic EZ-DM model.

**Figure 4.**
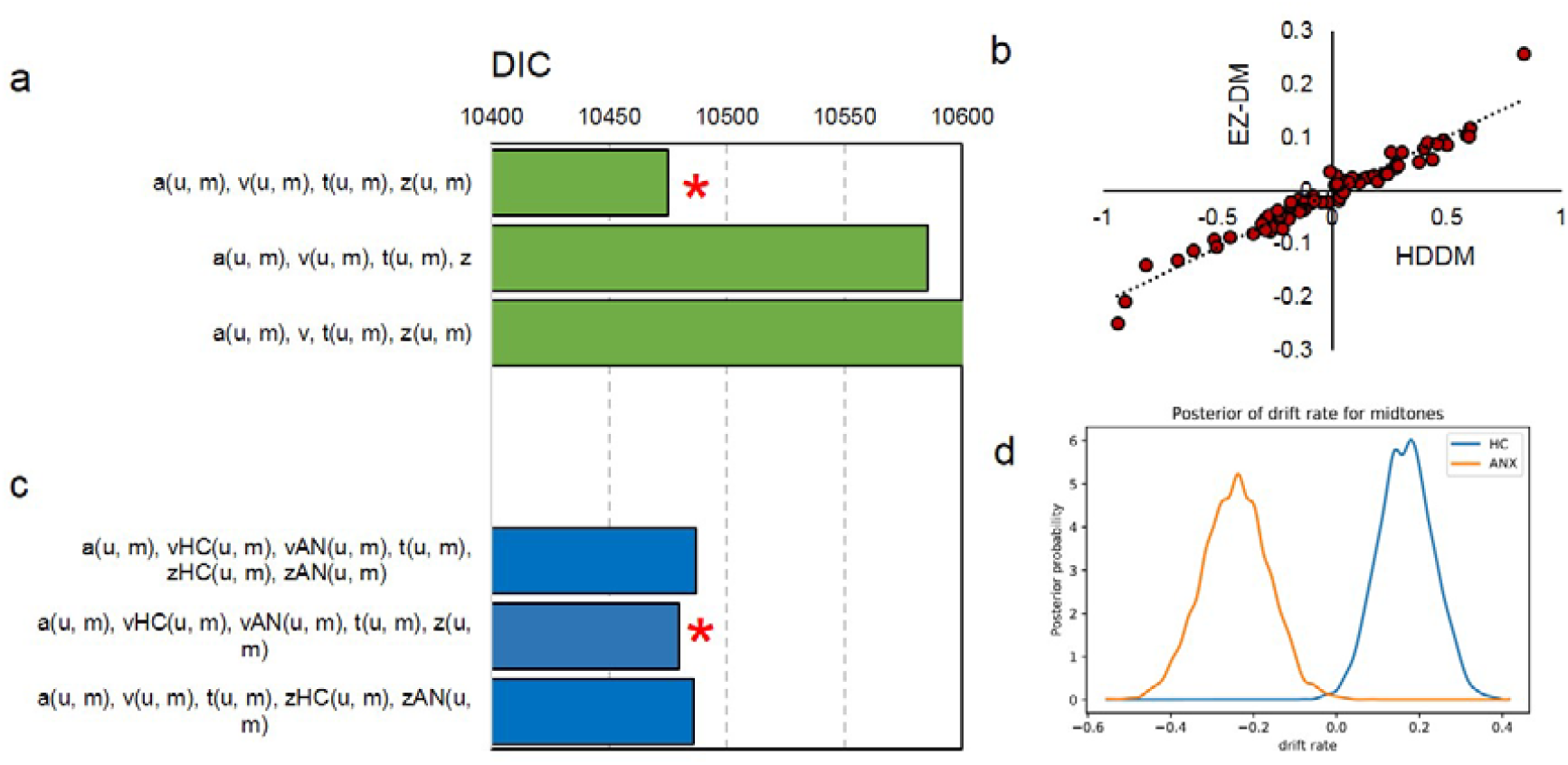
*hierarchical drift diffusion modelling of pathological anxiety* reveals **a**) a winning model (*) that includes separate drift rate (v), boundary separation (a) non-decision time (t) and bias (z) parameters for unambiguous (u) and ambiguous mid-tone (m) trial types based on lowest DIC scores. The v parameters recovered using this approach (HDDM) **b**) correlate tightly with those recovered from the EZ-DM model. Including group in the model fitting procedure **c**) demonstrates that the best model (*) fits the v parameter alone separately across groups. This is because, as can be seen on the posterior recovered samples, the **d**) v parameter was more negative in patients than controls. HC = asymptomatic Healthy Control; ANX = symptomatic individual

### Correlations

There was a strong positive correlation between affective bias and both drift rate measures (*r>*0.98, p*<*0.001), those who had a bias away from choosing high rewards had a slower drift rate towards high rewards.

There was weak evidence for a correlation between affective bias and STAI trait scores (*r* = −0.207, p*(two-tailed)* = 0.07, p*(one-tailed) =* 0.035) as well as weak evidence for a correlation between drift rate and STAI trait scores (EZDM *r* =−0.21, p*(two-tailed)* = 0.066 p*(one-tailed)* = 0.033; HDDM *r* =−0.22, p*(two-tailed)* = 0.053 p*(one-tailed)* = 0.027). In other words higher anxiety was associated with a reduced drift rate to the high reward choice. Additional exploratory correlations can be found in the supplement.

### Study 2

#### Threat of shock manipulation check

Participant anxiety ratings were significantly higher during the threat condition relative to the safe condition *t*_(44) =_ 8.92, p*<*0.001,*d =* 1.88 (safe mean = 1.64, SD = 1.05; threat mean = 4.93, SD = 2.21). Bayesian analysis provided decisive evidence that a model with a main effect of threat was the winning model (BF_10 =_ 4.68 e^8^).

#### Choice Behaviour

Accuracy for the high and low tones were high (Table 3) and comparable across conditions (*t*_*(46)*_ = 0.975, p = 0.335,*d* = 0.02, and *t*_*(46)*_ = 1.597, p = 0.117,*d =* 0.33, respectively). During the threat condition the proportion of mid-tones associated with high reward was smaller relative to the safe condition but did not achieve significance (*t*_(46) =_ 1.94, p = 0.06, *d =*−0.40; see Fig.3). Bayesian analysis anecdotally favoured a model with a main effect of condition (BF_10_ = 1.019).

**Table 3:** Average choice, accuracy, and reaction time (ms) to respond to tones in each condition in study 2.

| **Tone / Condition** | **Accuracy (standard deviation)** |
| --- | --- |
| Low tone (safe) | 0.99 (0.030) |
| High tone (safe) | 0.96 (0.047) |
| Low tone (threat) | 0.98 (0.036) |
| High tone (threat) | 0.95 (0.061) |
| **Condition** | **Proportion high reward responses to mid-tone** |
| Safe | 0.53 (0.20) |
| Threat | 0.49 (0.19) |
| **Tone / Condition** | **Reaction Time** |
| Low tone (safe) | 815.29 (192.09) |
| Low tone (threat) | 767.84 (206.84) |
| Mid-tone (safe) | 954.36 (185.63) |
| Mid-tone (threat) | 970.05 (168.97) |
| High tone (safe) | 830.46 (185.60) |
| High tone (threat) | 787.37 (205.78) |

#### Reaction time

See Table 3 for RT to different tone types across conditions. There was no difference between conditions in time taken to respond to the mid-tone (*t*_(46) =_ 1.24, p = 0.221,*d =* 0.26). Bayesian analysis confirmed that the null model was the winning model (BF_10_ = 0.325).

### DDM

#### EZ-DM

There was no significant difference between conditions in drift rate, non-decision time or boundary separation in decision-making to the mid-tones (*p*s> 0.125). Bayesian analysis confirmed that the null model was the winning model in all cases (BF_10_<1).

#### HDDM

The winning model comprised drift rate, boundary separation, non-decision time and bias parameter all fitted separately across ambiguous and unambiguous trials (figure 5a). This was the same model as study 1; however, this time, adding condition (figure 5b) into the hierarchy in this winning model resulted in substantially worse fits, thereby providing no justification for dividing trials by condition. In other words, the full hierarchical procedure again agreed with the EZ-DM procedure.

**Figure 5.**
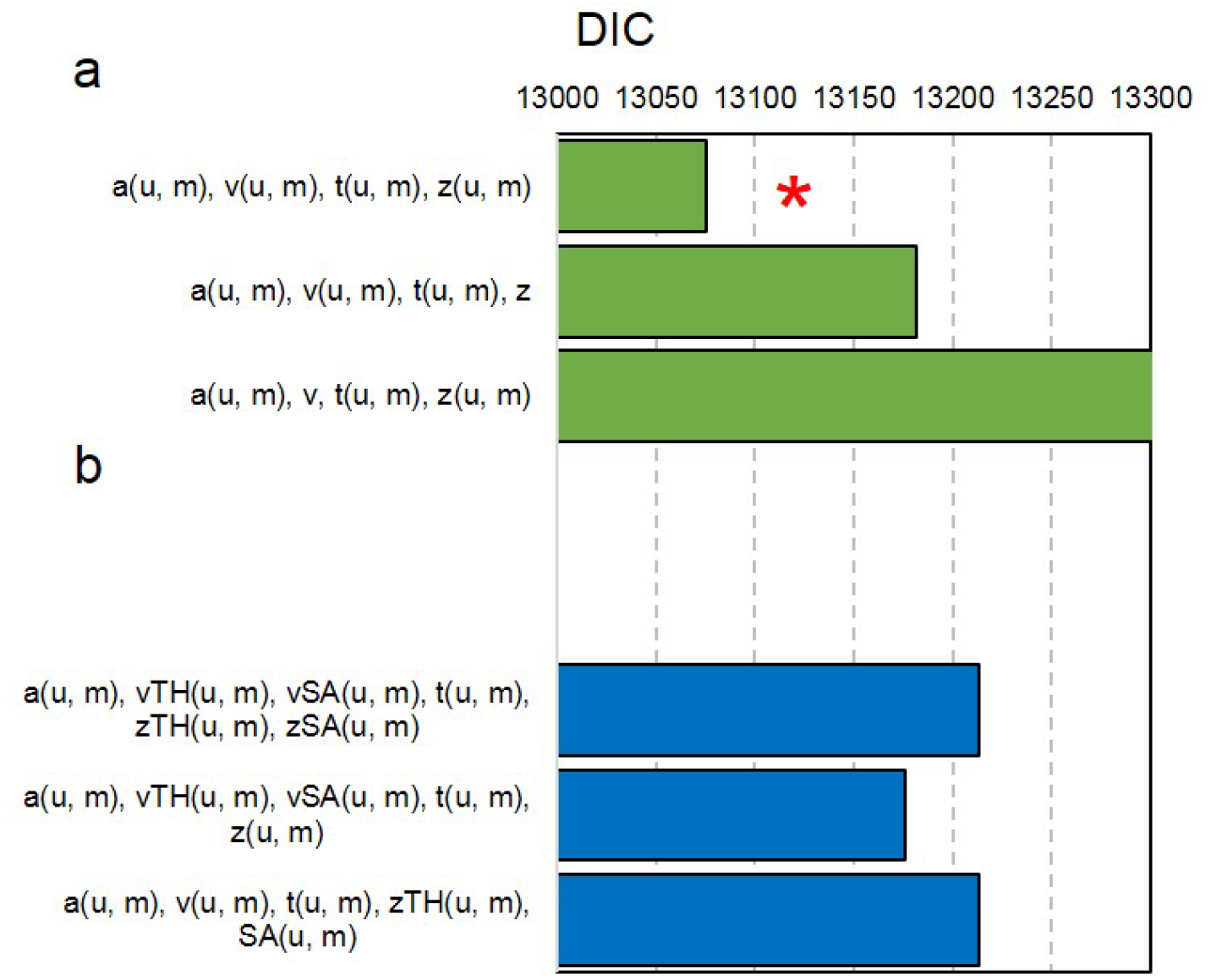
*hierarchical drift diffusion modelling of induced anxiety* reveals **a**) a winning model (*) that includes separate drift rate (v), boundary separation (a) non-decision time (t) and bias (z) parameters unambiguous (u) and ambiguous mid-tone (m) trial types based on lowest DIC scores. Including condition in the model fitting procedure **b**) provides substantially worse fits, thereby providing no evidence for an effect of condition.

## Conclusion

In this study we directly back-translate a rodent measure of affective bias. We demonstrate that pathological mood and anxiety disorders, but not transient induced anxiety in asymptomatic individuals, is associated with increased negative affective bias in task performance. This bias can, moreover, be attributed to reduced ‘drift rate’ on a computational model of reaction times.

Our results align with evidence documenting negative affective bias in mood and anxiety disorders ^4,22,37^ as well as two prior (conceptually different) studies^38,39^ linking mood disorder symptomatology to drift rates on the drift diffusion model. Critically, the anxiety-negative bias interaction translates the impact of a) acute anxiogenic pharmacological manipulation and b) chronic stress in the rodent task^15^ (Figure 6) into humans, suggesting that these rodent manipulations may be suitable preclinical screens for candidate therapeutics.

**Figure 6.**
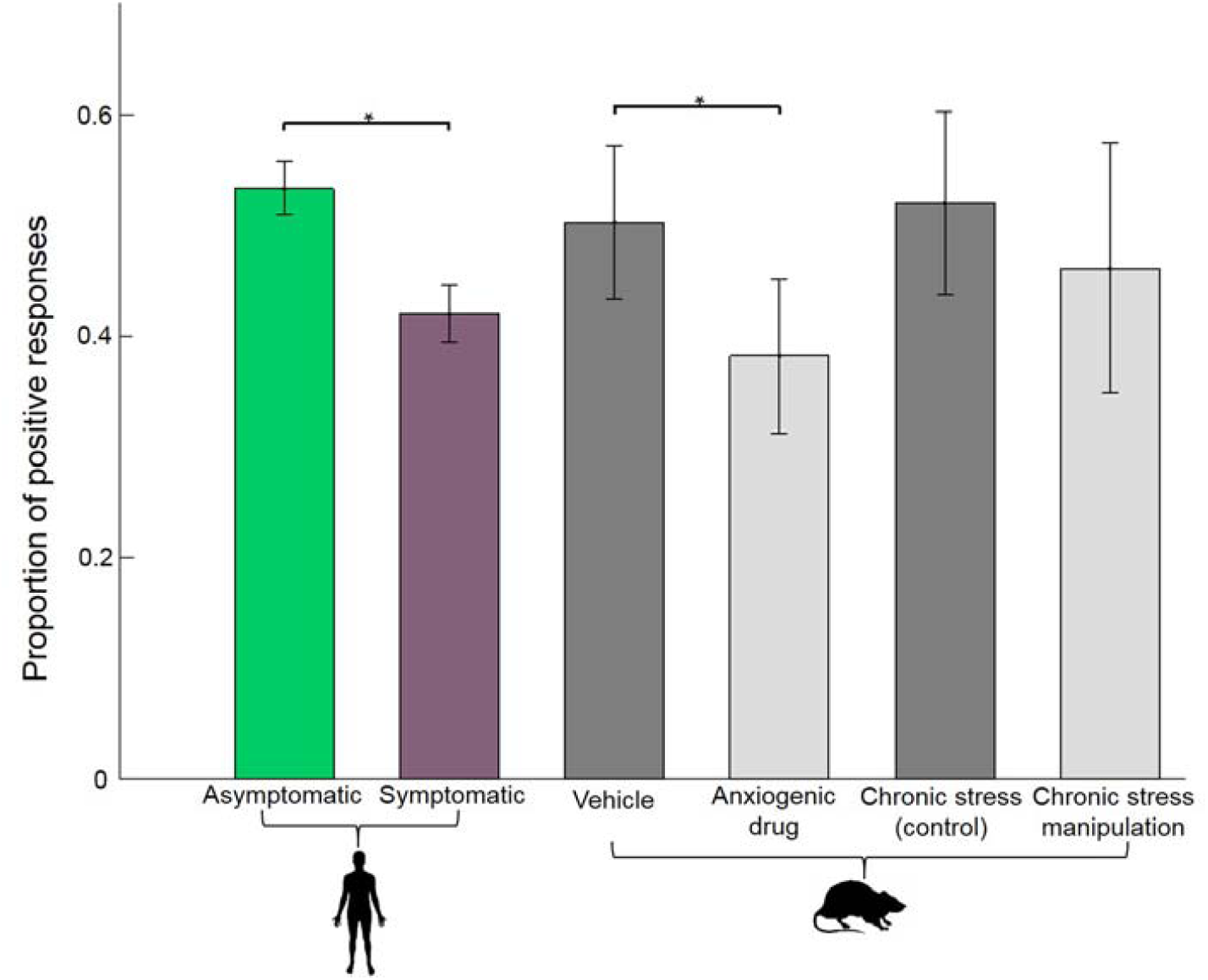
Cross-species performance comparison. Plots illustrating the overlap of human pathological anxiety and rodent anxiety models on choice performance. (\**p* < 0.05). Data presented in^15^. After acute pharmacological manipulation with FG7142 (3mg or 5mg; average dose plotted), rats showed an increased negative affective bias in choice behaviour on the ambiguous tone, relative to vehicle. For the chronic stress manipulation between weeks 3 and 4 post-stress intervention average of 6 post-stress intervention weeks plotted), rats showed an increased negative affective bias in choice behaviour on the ambiguous tone, relative to control.

Threat of shock instantiates negative affective biases across many areas of cognition^20^, but counter to predictions, induced anxiety in asymptomatic individuals did not reliably shift performance on this task. One potential explanation is that that, in the asymptomatic group, the induced anxiety task was always completed following the neutral version of the task. This may have increased familiarity with the task and counteracted any biases. However, it is also worth noting that the observation that decision-making is more sensitive to pathological than transient anxiety is also consistent with chronic vs acute restraint stress in rats^15^. Perhaps, therefore, acute environmental anxiety promotes *adaptive* harm-avoidance^20^, by increasing attentional and perceptual biases towards threats, without influencing higher-order decision-making processes. Supporting this is evidence demonstrating that, whilst encoding of values in ‘lower-level’ brain valuation structures changes as a function of threat-induced anxiety, decision-making behaviour remains unperturbed^40–42^ by threat of shock. It could therefore be that lower-level learning and memory are *immediately* influenced by transient states, but that the impact upon higher order processes builds up over time^43^. If correct, this suggests that, at least on the present measure, there is something quantifiably different between transient anxiety in healthy humans and pathological anxiety. From a clinical perspective this is unsurprising, but it is notable because some effects do overlap across induced and pathological anxiety^20,23,44^. Finally, it is worth acknowledging that we may simply be underpowered to detect an effect of threat, perhaps because the manipulation was not strong enough. This is arguably unlikely considering increased anxiety ratings under threat, and the wide-ranging influence of induced anxiety on cognition^20^. However, if correct it would mean that the within-subject effect of transient anxiety is considerably smaller than the between-subject effect detected in the group study.

The dissociation across studies suggests, however, that this task is more sensitive to the pathological state than transient changes in anxiety in asymptomatic individuals. On reflection, this extends the translational potential of this paradigm. Given the failure of many preclinical to phase 1 clinical trials^10,11^, translational paradigms which are *more* sensitive to the clinical state than transient mood changes are valuable. Modifying affective biases in mood and anxiety disorders is crucial given their proposed role in the development and maintenance of symptoms^5 – 7^. Both pharmacological and psychological treatments^45 – 47^, are thought to exert their effects via altering affective biases^5^. In rodents, for instance, a similar task has been shown to be sensitive to anxiolytic manipulations; a positive bias is exhibited after treatment with the antidepressant Venlafaxine^48^. Confirming the same effect on this task in a medicated human sample would therefore enhance the predictive validity of this task for drug testing new anxiolytics.

In addition to facilitating screening of novel anxiolytics, the present translational pipeline provides a potential means of understanding the mechanisms underpinning this negative bias^49^. Running causal studies in rodents can help us delineate the neurobiological processes underpinning biased choices on this task^12^. Moreover, linking task performance to a formal model of decision-making (DDM) provides a step towards bridging the gap between brain and behaviour. Notably, in the rodent model that most closely mimics the behaviour of anxious humans^15^, as well as in humans with anxiety disorders, the drift rate parameter was reduced. This suggests that, in both cases, anxiety reduces the rate of evidence accumulation (although it should be noted that, unlike the rodent mode, in humans the bias parameter was unaffected). Crucially, the parameters of this model are thought to be biophysically plausible; they can be computed by populations of neurons^50^; taking us closer to being able to link underlying neural activity to psychiatric symptoms. Such links are necessary for a full mechanistic account of psychiatric symptoms and are the guiding principal of the burgeoning field of computational psychiatry^51^.

Ultimately, we argue that improved treatments are unlikely without a better understanding of the underlying biological mechanisms that any putative treatments are attempting to target. Given the huge costs of mood and anxiety disorders; as well as the large number of individuals for whom none of our current treatments work; new and improved treatments, and therefore better methods of screening for such treatments, are long overdue. We propose that the task presented here may hold promise as a means of better screening for candidate treatments across humans and animal models.

## Acknowledgements

This research was funded by A Medical Research Foundation Equipment Competition Grant (C0497, Principal Investigator OJR), and a Medical Research Council Career Development Award to OJR (MR/K024280/1).

## Author Contributions

O.J.R. conceived of the experiment. E.R conceived of the original animal task and provided detailed comments on the interpretation. O.J.R. wrote the task scripts with guidance from C.H. J.A completed testing and data collection. J.A completed data analysis under the supervision of O.J.R. J.A. and O.J.R wrote first draft of the paper and C.H. and E.R. provided critical feedback. All authors approved the final version for submission.

## References

1 Beddington J, Cooper CL, Field J, et al. The mental wealth of nations. Nature 2008; 455: 1057–60.

2 MacLeod C, Mathews A, Tata P. Attentional bias in emotional disorders. J Abnorm Psychol 1986; 95: 15–20.

3 Mogg K, Bradley BP. Time course of attentional bias for fear-relevant pictures in spider-fearful individuals. Behav Res Ther 2006; 44: 1241–50.

4 Hirsch C, Mathews A. Interpretative inferences when reading about emotional events. Behav Res Ther 1997; 35: 1123–32.

5 Roiser JP, Elliott R, Sahakian BJ. Cognitive Mechanisms of Treatment in Depression. Neuropsychopharmacology 2012; 37: 117–36.

6 Kendler KS, Kuhn J, Prescott CA. The interrelationship of neuroticism, sex, and stressful life events in the prediction of episodes of major depression. Am J Psychiatry 2004; 161: 631–6.

7 Harmer CJ, Goodwin GM, Cowen PJ. Why do antidepressants take so long to work? A cognitive neuropsychological model of antidepressant drug action. Br J Psychiatry 2009; 195: 102–8.

8 Psychological Therapies: Annual report on the use of IAPT services Psychological Therapies: Annual Report on the use of IAPT services, England, 2015-16. 2016.

9 Joffe RT, Levitt AJ, Sokolov ST. Augmentation strategies: focus on anxiolytics. J Clin Psychiatry 1996; 57 Suppl 7: 25-31; discussion 32-3.

10 Scannell JW, Bosley J, Kyriakopoulou A, Serghiou S, Wilde A de, Sherratt N. When Quality Beats Quantity: Decision Theory, Drug Discovery, and the Reproducibility Crisis. PLoS One 2016; 11: e0147215.

11 Choi DW, Armitage R, Brady LS, et al. Medicines for the Mind: Policy-Based “Pull” Incentives for Creating Breakthrough CNS Drugs. Neuron 2014; 84: 554–63.

12 Badre D, Frank MJ, Moore CI. Interactionist Neuroscience. Neuron 2015; 88: 855–60.

13 Porsolt RD, Bertin A, Jalfre M. Behavioral despair in mice: a primary screening test for antidepressants. Arch Int Pharmacodyn Ther 1977; 229: 327–36.

14 Steru L, Chermat R, Thierry B, Simon P. The tail suspension test: a new method for screening antidepressants in mice. Psychopharmacology (Berl) 1985; 85: 367–70.

15 Hales CA, Robinson ESJ, Houghton CJ, Gotlib I, Mathews A, Spanagel R. Diffusion Modelling Reveals the Decision Making Processes Underlying Negative Judgement Bias in Rats. PLoS One 2016; 11: e0152592.

16 Robinson OJ, Letkiewicz AM, Overstreet C, Ernst M, Grillon C. The effect of induced anxiety on cognition: threat of shock enhances aversive processing in healthy individuals. Cogn Affect Behav Neurosci 2011; 11: 217–27.

17 Aylward J, Robinson OJ. Towards an emotional ‘stress test’: a reliable, non-subjective cognitive measure of anxious responding. Sci Rep 2017; 7: 40094.

18 Boureau Y-L, Dayan P. Opponency revisited: competition and cooperation between dopamine and serotonin. Neuropsychopharmacology 2011; 36: 74–97.

19 Robinson OJ, Krimsky M, Grillon C. The impact of induced anxiety on response inhibition. Front Hum Neurosci 2013; 7: 69.

20 Robinson OJ, Vytal K, Cornwell BR, Grillon C. The impact of anxiety upon cognition: perspectives from human threat of shock studies. Front Hum Neurosci 2013; 7: 203.

21 Robinson OJ, Charney DR, Overstreet C, Vytal K, Grillon C. The adaptive threat bias in anxiety: Amygdala–dorsomedial prefrontal cortex coupling and aversive amplification. Neuroimage 2012; 60: 523–9.

22 Anderson MH, Hardcastle C, Munafò MR, Robinson ESJ. Evaluation of a novel translational task for assessing emotional biases in different species. Cogn Affect Behav Neurosci 2012; 12: 373–81.

23 Robinson OJ, Chase HW. Learning and Choice in Mood Disorders: Searching for the Computational Parameters of Anhedonia. Comput Psychiatry 2017; 1: 208–33.

24 Ratcliff R, Smith PL, Brown SD, McKoon G. Diffusion Decision Model: Current Issues and History. Trends Cogn Sci 2016; 20: 260–81.

25 Wagenmakers E-J, Van Der Maas HLJ, Grasman RPPP. An EZ-diffusion model for response time and accuracy. Psychon Bull Rev 2007; 14: 3–22.

26 van Ravenzwaaij D, Donkin C, Vandekerckhove J. The EZ diffusion model provides a powerful test of simple empirical effects. Psychon Bull Rev 2016;: 1–10.

27 Wiecki T V, Sofer I, Frank MJ. HDDM: Hierarchical Bayesian estimation of the Drift-Diffusion Model in Python. Front Neuroinform 2013; 7: 14.

28 MacLeod C, Mathews A. Cognitive bias modification approaches to anxiety. Annu Rev Clin Psychol 2012; 8: 189–217.

29 Lecrubier Y, Sheehan D, Weiller E, et al. The Mini International Neuropsychiatric Interview (MINI). A short diagnostic structured interview: reliability and validity according to the CIDI. Eur Psychiatry 1997; 12: 224–31.

30 Spielberger CD, Gorsuch LR, Lushene RE, Vagg PR, Jacobs GA. Manual for the State-Trait Anxiety Inventory. Palo Alto, CA: Consulting Psychologists Press, 1983.

31 Schmitz A, Grillon C. Assessing fear and anxiety in humans using the threat of predictable and unpredictable aversive events (the NPU-threat test). Nat Protoc 2012; 7: 527–32.

32 Grillon C, Ameli R, Merikangas K, Woods SW, Davis M. Measuring the time course of anticipatory anxiety using the fear-potentiated startle reflex. Psychophysiology 1993; 30: 340–6.

33 Grillon C, Baas JMP, Cornwell B, Johnson L. Context Conditioning and Behavioral Avoidance in a Virtual Reality Environment: Effect of Predictability. Biol Psychiatry 2006; 60: 752–9.

34 JASP. JASP (Version 0.7.5.5). 2016.

35 Jeffreys H. The theory of probability. 1998.

36 Patil A, Huard D, Fonnesbeck CJ. PyMC: Bayesian Stochastic Modelling in Python. J Stat Softw 2010; 35: 1–81.

37 Mathews A. Effects of modifying the interpretation of emotional ambiguity. J Cogn Psychol 2012; 24: 92–105.

38 White CN, Ratcliff R, Vasey MW, McKoon G. Anxiety enhances threat processing without competition among multiple inputs: A diffusion model analysis. Emotion 2010; 10: 662–77.

39 Dillon DG, Wiecki T, Pechtel P, et al. A computational analysis of flanker interference in depression. Psychol Med 2015; 45: 2333–44.

40 Engelmann JB, Meyer F, Fehr E, Ruff CC. Anticipatory Anxiety Disrupts Neural Valuation during Risky Choice. J Neurosci 2015; 35.

41 Charpentier CJ, Hindocha C, Roiser JP, Robinson OJ, Iverson G. Anxiety promotes memory for mood-congruent faces but does not alter loss aversion. Sci Rep 2016; 6: 24746.

42 Robinson, O.J., Bond RL, Roiser JP. The impact of threat of shock on the framing effect and temporal discounting: executive functions unperturbed by acute stress? Front Psychol 2015; 6: 1315.

43 Anderson MH, Munafò MR, Robinson ESJ. Investigating the psychopharmacology of cognitive affective bias in rats using an affective tone discrimination task. Psychopharmacology (Berl) 2013; 226: 601–13.

44 Robinson OJ, Krimsky M, Lieberman L, Allen P, Vytal K, Grillon C. The dorsal medial prefrontal (anterior cingulate) cortex–amygdala aversive amplification circuit in unmedicated generalised and social anxiety disorders: an observational study. The Lancet Psychiatry 2014; 1: 294–302.

45 Fournier JC, DeRubeis RJ, Hollon SD, et al. Antidepressant Drug Effects and Depression Severity. JAMA 2010; 303: 47.

46 Zarate CA, Singh JB, Carlson PJ, et al. A Randomized Trial of an N-methyl-D-aspartate Antagonist in Treatment-Resistant Major Depression. Arch Gen Psychiatry 2006; 63: 856.

47 Dimidjian S, Hollon SD, Dobson KS, et al. Randomized trial of behavioral activation, cognitive therapy, and antidepressant medication in the acute treatment of adults with major depression. J Consult Clin Psychol 2006; 74: 658–70.

48 Hinchcliffe JK, Stuart SA, Mendl M, Robinson ESJ. Further validation of the affective bias test for predicting antidepressant and pro-depressant risk: effects of pharmacological and social manipulations in male and female rats. Psychopharmacology (Berl) 2017; published online July. DOI:10.1007/s00213-017-4687-5.

49 Stuart SA, Butler P, Munafò MR, Nutt DJ, Robinson ES. Distinct Neuropsychological Mechanisms May Explain Delayed-Versus Rapid-Onset Antidepressant Efficacy. Neuropsychopharmacology 2015; 40: 2165–74.

50 Ratcliff R, McKoon G. The diffusion decision model: theory and data for two-choice decision tasks. Neural Comput 2008; 20: 873–922.

51 Huys QJM, Maia T V, Frank MJ. Computational psychiatry as a bridge from neuroscience to clinical applications. Nat Neurosci 2016; 19: 404–13.

